# A chromosomal inversion acts as a hotspot for divergence in multiple ecologically relevant traits

**DOI:** 10.64898/2026.10.08.757055

**Authors:** Juliana Rodríguez-Fuentes, Lynn Wedekind, Lea Stadelmann, Nicole Nesvadba, Verena Saladin, Marius Roesti, Catherine L. Peichel

**Affiliations:** Division of Evolutionary Ecology, Institute of Ecology and Evolution, University of Bern, Bern 3012, Switzerland

**Keywords:** Chromosomal inversion, Phenotypic divergence, Local adaptation, *Gasterosteus aculeatus*

## Abstract

Chromosomal inversions are proposed to aid local adaptation to divergent habitats when there is gene flow by creating regions of low recombination that can maintain tight linkage between multiple locally adaptive alleles. However, key theoretical predictions about the role of chromosomal inversions in local adaptation remain largely untested in empirical systems. Here, we test the prediction that inversions will concentrate relatively small genetic effects across multiple ecologically relevant traits by exploring inversion–phenotype associations of three inversions in threespine stickleback (*Gasterosteus aculeatus*). Using complementary approaches of association mapping in wild-caught individuals and genetic mapping in lab-raised individuals from two divergent genomic backgrounds, we found that an inversion on chromosome XXI influences multiple ecologically relevant traits. The inversion has small-to-moderate effects on these traits, as expected for inversions involved in polygenic adaptation. Further consistent with theoretical predictions, inversion heterozygotes show intermediate phenotypes. While most inversion–phenotype associations are consistent across different genomic backgrounds and experimental approaches, some inversion–phenotype associations were specific to a single dataset. These results indicate that the phenotypic effects of inversions are shaped by both genomic and environmental context, which could facilitate population-specific adaptation. Together, these results support the role of the chromosome XXI inversion in ecological divergence in threespine stickleback. More broadly, our results suggest that inversions can contribute to local adaptation by preserving linkage between small-effect adaptive alleles despite extensive gene flow.

## Introduction

Adaptation to divergent environments commonly requires the evolution of differences in multiple traits that together increase fitness. However, in the presence of maladaptive gene flow, recombination can mix locally adaptive with foreign genetic variation, thereby disrupting beneficial associations among loci underlying adaptive traits. Thus, gene flow can break apart adaptive trait combinations generated by divergent selection (Felsenstein, 1981). Theoretical work has proposed that divergence can nevertheless proceed through mechanisms that create or maintain linkage among loci underlying adaptive traits (Bürger & Akerman, 2011; Yeaman & Whitlock, 2011).

Chromosomal inversions may provide such a mechanism for protecting adaptive trait combinations through their effects on recombination (Faria et al., 2019; Hoffmann & Rieseberg, 2008; Schluter & Rieseberg, 2022; Wellenreuther & Bernatchez, 2018). Inversions are a type of structural mutation in which the orientation of a DNA segment within a chromosome is reversed relative to the standard orientation. In inversion heterozygotes, crossing over between the different inversion orientations can produce unbalanced recombinant chromosomes that result in non-viable offspring. The loss of these recombinant haplotypes effectively suppresses recombination between the standard and inverted orientations. Inversions can therefore facilitate population divergence by maintaining different combinations of alleles underlying multiple adaptive traits in alternative orientations (Hoffmann & Rieseberg, 2008; Kirkpatrick, 2010; Sturtevant & Beadle, 1936). When these orientations are favored in different habitats and heterozygotes have reduced fitness because they express phenotypes that are not adaptive in either habitat, selection can maintain divergence despite ongoing gene flow by keeping divergent alleles in the alternative arrangements (Kirkpatrick & Barton, 2006). Inversions are particularly predicted to contribute to phenotypic divergence and local adaptation when adaptive alleles are prone to being lost through gene flow; that is, when adaptive trait divergence is polygenic and individual adaptive traits are influenced by loci of relatively small effects (Roesti, 2018; Schaal et al., 2022). Taken together, theory predicts that when inversions facilitate local adaptation through their effect on suppressing recombination, multiple traits will be influenced by the inversion, with individual alleles within the inversion having relatively small effects on these traits, and that heterozygotes will have phenotypes that differ from individuals that are homozygous for either of the alternative orientations.

Consistent with these theoretical predictions, empirical studies of ecotypes or closely related species from divergent habitats often find strong frequency differences of alternative inversion orientations (Wellenreuther & Bernatchez, 2018). In some cases, inversions have further been shown to influence multiple traits, often with relatively small to moderate effect sizes and intermediate phenotypes in heterozygotes (Akopyan et al., 2026; Hager et al., 2022; Kapun et al., 2016; Koch et al., 2021; Lowry & Willis, 2010; Sanchez-Donoso et al., 2022). However, some studies investigating the phenotypic effects of inversions have relied on quantitative trait locus (QTL) mapping in crosses between divergent populations or ecotypes. In such crosses, inversion genotypes can remain strongly linked to the neighboring genomic background inherited from the parental populations, making it difficult to distinguish phenotypic effects of the inversion itself from those of linked genetic variation. Genome-wide association studies (GWAS) can provide finer mapping resolution by exploiting historical recombination in natural populations but require suitable populations in which both the inversion and relevant phenotypic variation segregate. Moreover, because both QTL mapping and GWAS are typically phenotype-driven, they are not necessarily well suited to identifying the range of traits associated with a particular inversion of interest.

An alternative approach is therefore to start from a naturally polymorphic population with a known segregating inversion and systematically identify the phenotypic traits associated with different inversion genotypes (e.g. Sanchez-Donoso et al., 2022). In such a mapping population, historical recombination is expected to have reduced linkage between the inversion and its neighboring genomic background, allowing its phenotypic associations to be resolved more precisely. This inversion association-based mapping approach can be complemented with controlled crosses where individuals from the same population segregate for the alternative inversion genotypes against a shared genomic background (e.g. Kapun et al., 2016). Raising these offspring under common environmental conditions allows the genetic component of inversion-associated phenotypic differences to be assessed independently of environmental variation. Together, these complementary approaches can reveal the range and magnitude of phenotypic effects associated with an inversion in nature, test whether these effects have a genetic basis under controlled conditions, and characterize the phenotypes of heterozygotes relative to both homozygous orientations. They therefore provide a powerful framework for testing how inversions contribute to multi-trait adaptive divergence.

Threespine stickleback fish (*Gasterosteus aculeatus*) offer a great system for applying this framework to test predictions about how inversions contribute to local adaptation through their effects on recombination. This northern-temperate fish is an ancestrally marine species that repeatedly colonized newly formed freshwater habitats following glacier retreat after the last Ice Age (Bell & Foster, 1994; McKinnon & Rundle, 2002). Marine and freshwater sticklebacks have diverged in many morphological, behavioral, and physiological traits, often resulting in repeated (or parallel) phenotypic syndromes that differentiate freshwater and marine ecotypes (Bell & Foster, 1994). For example, marine fish have strong bony armor, composed of a complete set of bony lateral plates and long spines, as well as longer and more numerous gill rakers (Bell & Foster, 1994). Freshwater fish that live in benthic habitats often have reduced armor and shorter and fewer gill rakers than marine fish (Bell & Foster, 1994). This contrasts with freshwater fish from limnetic habitats, which are more heavily armored than benthic sticklebacks and have gill raker phenotypes similar to those of marine sticklebacks (Gross & Anderson, 1984; Kaeuffer et al., 2012; McPhail, 1992; Roesti et al., 2023; Schluter & McPhail, 1992; Vamosi & Schluter, 2004). Similarly, the lateral line sensory system of limnetic fish is more similar to that of marine than benthic fish (Wark & Peichel, 2010). Thus, phenotypic divergence in sticklebacks is not restricted to the marine–freshwater axis but also occurs among freshwater ecotypes (e.g., Berner et al., 2008; Deagle et al., 2012; Roesti et al., 2023; Schluter & McPhail, 1992; Stuart et al., 2017; Willacker et al., 2010). Furthermore, divergence in most traits has a polygenic genetic architecture (Peichel & Marques, 2017). This extensive knowledge of genetic and phenotypic divergence across multiple ecologically relevant traits between different stickleback ecotypes provides an excellent opportunity to test whether chromosomal inversions keep adaptive trait combinations together, as predicted if they contribute to local adaptation.

Consistent with the presence of gene flow among stickleback ecotypes (Fang et al., 2020; Haenel et al., 2019; Hagen, 1967; Schluter & Conte, 2009), three chromosomal inversions, located on chromosomes (chr) I, XI, and XXI are proposed to contribute to local adaptation in threespine sticklebacks. These are relatively small inversions in the 450 megabase (Mb) stickleback genome; the chrI and chrXI inversions are under 0.5 Mb, and the chrXXI inversion is approximately 1.7 Mb. Across these regions, they show strong genomic divergence between marine and freshwater populations from both the Atlantic and Pacific basins (Adhikari et al., 2025; Fang et al., 2020; Jones et al., 2012; Magalhaes et al., 2021; Roberts Kingman et al., 2021; Terekhanova et al., 2019) and can also differ in frequency among freshwater ecotypes (Brachmann et al., 2026; Haenel et al., 2019; Magalhaes et al., 2021; Marques et al., 2022; Roesti et al., 2015). Here, we will refer to the alternative inversion orientations as “marine” or “freshwater” based on where they were first identified (Jones et al., 2012). Importantly, these inversions are ancient, and their freshwater orientations persist as standing genetic variation in marine populations across much of the species’ geographic range (Jones et al., 2012; Nelson & Cresko, 2018). This widespread availability as pre-existing variation may allow selection to repeatedly act on these inversions following freshwater colonization from the sea, facilitating their repeated use in adaptation (Fang et al., 2020; Nelson & Cresko, 2018). However, whether these inversions are associated with multiple divergent traits has not been directly tested. Although a meta-analysis indicated that the chrXXI inversion harbors more QTL than expected by chance (Peichel & Marques, 2017), the QTL regions are large and their phenotypic effects cannot be attributed directly to the inversion (Erickson et al., 2016). Indeed, a QTL explaining tooth morphology was found to overlap the chrXXI inversion, but a follow-up study showed that the causative locus is outside of the inversion (Cleves et al., 2014). Thus, it remains unclear whether the inversions themselves harbor loci that underlie multiple adaptive traits.

Here, we tested three predictions about the contribution of the chrI, XI, and XXI inversions to adaptive phenotypic divergence: (1) the inversions influence multiple ecologically relevant traits; (2) their effects on individual traits are relatively small; and (3) inversion heterozygotes differ phenotypically from homozygous individuals. To test these predictions, we used two complementary approaches. First, we performed association mapping in wild fish from an Atlantic stream population, which is polymorphic at the three inversions (Roesti et al., 2015). Second, we generated laboratory-reared crosses segregating the alternative inversion orientations on a common genomic background. One set of crosses used Atlantic stream fish from the same population used for association mapping, and the other used marine fish from the Pacific basin. Together, these approaches allowed us not only to test the predicted phenotypic effects of the inversions, but also to assess whether inversion–phenotype associations are consistent across genomic backgrounds, environments, and experimental approaches.

## Materials and methods

### Fish collection and crosses

#### Wild population for inversion-based association mapping

For association mapping in a wild population, sticklebacks were collected using unbaited minnow traps from a naturally inversion-polymorphic stream population in the Lake Constance watershed: Riedgraben in Grasbeuren, Germany (GRA; 47.71909, 9.31563). This population is polymorphic for the chrI, XI, and XXI inversions (Roesti et al., 2015). Fish were collected during June in three sampling years: 2020 (n = 221), 2021 (n = 28), and 2024 (n = 584). Individuals were euthanized using a lethal dose of buffered MS-222 (0.2% tricaine methane sulfonate), blotted dry, and weighed using a digital scale with a precision of 0.01 mg. The right pectoral fin was removed and stored in 100% ethanol, and the body was stored in freshly made 4% paraformaldehyde (PFA) at 4°C for 24-36 hours before further processing (see Phenotyping: Assessing variation across different trait sets). Fish sampling was conducted under a permit issued by the local fisheries office.

#### Laboratory crosses from the Atlantic stream population

For testing inversion-associated phenotypic differences under controlled conditions in a freshwater genomic background, one set of laboratory-raised individuals was derived from fish collected from the GRA stream population in 2021. First-generation (F1) crosses were generated in the field using in-vitro fertilization as described in Roesti et al (2026). These F1 fish were genotyped at the chrI, XI, and XXI inversions, and individuals carrying both the marine and freshwater orientations (i.e., inversion heterozygotes) were identified. Only F1 heterozygotes at the chrXXI inversion were crossed to generate second-generation (F2) crosses, which were raised to adulthood. Both F1 and F2 crosses were reared under standardized conditions as described in Roesti et al (2026). Adult fish from three F2 crosses were sacrificed using a lethal dose of buffered MS-222. The right pectoral fin was removed and stored in 100% ethanol, and the bodies were stored in freshly made 4% PFA at 4°C for 24–36 hours before further processing (see Phenotyping: Assessing variation across different trait sets).

#### Laboratory crosses from a Pacific marine population

To test inversion-associated phenotypic differences in a marine genomic background, another set of laboratory-raised individuals consisted of F1 crosses derived from wild-caught inversion heterozygotes from an eastern Pacific Ocean population (Oyster Lagoon, British Columbia, Canada; 49.612056, -124.031194), as described in Rodríguez-Fuentes et al. (2026). F1 crosses were generated between fish heterozygous at a single inversion and homozygous for the marine orientation at the other two inversions. Six independent F1 crosses were generated for each inversion. Individuals were reared under standardized conditions for 14–16 months and euthanized with a lethal dose of buffered MS-222. The right pectoral fin was removed and stored in 100% ethanol, while the bodies were either stored in freshly made 4% PFA at 4°C for 24–36 hours or 10% formalin for one week, depending on the phenotypes of interest (see Phenotyping: Assessing variation across different trait sets). Protocols for animal husbandry and crosses were approved by the relevant authorities.

### Inversion genotyping and sexing

Fish were genotyped at the inversions and for sex using DNA extracted from fin clips stored in 100% ethanol, as described in Rodríguez-Fuentes et al. (2026). Briefly, DNA was extracted from a fin clip using a modified HotSHOT protocol (Archambeault et al., 2020; Meeker et al., 2007). Extracted DNA was used as a template for PCR amplification of diagnostic indels that distinguish between the alternative inversion orientations and between the X and Y sex chromosomes (Rodríguez-Fuentes et al., 2026). At the chromosomal inversions, individuals can be homozygous for the marine-(MAR) or freshwater (FW)-orientation or heterozygous (HET).

### Phenotyping: Assessing variation across different trait sets

Wild-caught (Table S1) and lab-raised individuals (Table S2) were phenotyped for different sets of morphological traits, including external count traits (lateral line neuromast and lateral plate number), linear body shape traits, and branchial feeding traits. 406 wild-caught individuals from the Atlantic-derived GRA stream population were phenotyped for all traits. These included all individuals sampled in 2020 and 2021 and a subset from 2024, added to increase the sample sizes of FW and HET individuals for the chrXI and XXI inversions. From the lab-raised GRA stream crosses, 248 individuals were phenotyped for the external count traits and linear body shape traits. From the lab-raised marine crosses, 806 individuals were phenotyped: 285 for external count traits, 419 for linear body shape and branchial feeding traits, and 102 for all traits. All individuals that were phenotyped for all traits were first stained to highlight neuromasts (see External count traits: lateral line neuromasts and plates) and then stained to highlight bony tissue (see Linear body shape traits and Branchial feeding traits).

#### External count traits: lateral line neuromasts and plates

Individuals were phenotyped for the number of neuromasts in twelve lines of the lateral line system (Fig. S1C, Wark et al., 2012; Wark & Peichel, 2010). To visualize neuromasts, individuals were stained using alkaline phosphatase. Before staining, individuals stored in 4% PFA were washed with phosphate-buffered saline (PBS) and stored in this solution at 4°C. Alkaline phosphatase staining followed the protocol described in Archambeault et al. (2020). Neuromasts in each line and lateral bony plates were counted on the left side of stained fish under a Nikon SMZ1500 stereomicroscope (Nikon Instruments, Tokyo, Japan). Since staining quality may affect neuromast counts, staining quality was scored for each fish on a scale of 1–4. Measurements and staining quality assessment were performed by a single observer per dataset, except for the lab-raised marine dataset, which was measured by two observers. However, all individuals within a given cross were scored by the same observer.

#### Linear body shape traits

Individuals were stained with Alizarin Red to highlight bony tissue and phenotyped for a set of linear body shape traits. Individuals were fixed in 10% formalin for at least one week, after which the formalin was removed through multiple washes with water. Individuals were then placed in the staining solution (1 g/L Alizarin Red, 0.01 g/L KOH) for two to three days depending on staining progression. For fish stained for three days, the staining solution was replaced on the second day. Staining progression was checked under a Nikon SMZ1500 stereomicroscope (Nikon Instruments, Tokyo, Japan), and staining was stopped by multiple washes with water. Alizarin Red-stained individuals were stored in 37% isopropyl alcohol until phenotyping.

Linear body shape was characterized from photographs of Alizarin Red-stained fish using a set of linear measurements (“traits”). Photographs were taken from the left side of pinned individuals, with a ruler and individual tag included in each image, using a Nikon D850 camera (Nikon Instruments, Tokyo, Japan). Linear body shape traits were derived from a set of 30 landmarks established by Roesti et al. (2026) that capture morphological variation among ecotypes related to defense, feeding, vision, and swimming (Fig. S1A; Table S3).

Landmarks were placed automatically using a machine learning approach using the simple-ML-morph pipeline (Lürig, 2022; Porto & Voje, 2020; Sulser et al., 2026). Automated landmarking was performed independently for wild-caught and lab-raised datasets because a single model could not accurately predict landmarks across datasets. Models were trained on randomly selected subsets of photographs that were manually landmarked using tpsDIG v2.32 (Rohlf, 2015), including 190 wild-caught stream, 51 lab-raised stream, and 200 lab-raised marine individuals. Because the original training set for the lab-raised stream dataset did not produce sufficiently accurate landmarks, 85 additional photographs from the already automatically landmarked lab-raised marine dataset were added to the lab-raised stream training set (n = 136).

Each subset of manually landmarked photographs was split randomly into 80% training data and 20% test data. Different models were trained by varying the parameters *threads*, *tree depth*, and *cascade depth*, while keeping the remaining parameters at their default values. Model performance was compared using the pixel distance between the landmarks from the training and test data. The model with the lowest pixel distance between predicted and manually placed landmarks in the training and test data was to predict landmarks in the complete dataset. Model parameters for wild-caught stream and lab-raised marine datasets were *threads* = 12, *tree depth* = 2, *cascade depth* = 35, *oversampling* = 300, *feature pool size* = 1200; model parameters for lab-raised stream were: *threads* = 12, *tree depth* = 2, *cascade depth* = 50, *oversampling* = 300, *feature pool size* = 500). A single observer per dataset inspected and adjusted the predicted landmarks. Two additional landmarks were placed manually by a single observer to serve as a scale on the wild-caught stream and lab-raised marine datasets. On the lab-raised stream dataset, the scale distance was determined using a custom script to measure automatically the ruler (Sulser et al., 2026). Linear measurements were calculated as the pixel distance between sets of two landmarks (Table S3) using the function *LinearDist* (Adams & Otárola-Castillo, 2013). Pixel distances were converted to mm using the pixel distance from the scale. This resulted in 18 measurements, which were reduced to 17 linear traits by defining eye diameter as the average of the horizontal and vertical eye measurements.

#### Branchial feeding traits

Branchial traits (gill arch length, gill raker number, gill raker length) were measured on the first branchial arch of lab-raised marine and wild-caught stream individuals using a modified protocol from Berner et al. (2009) and Roesti et al. (2023) (Fig. S1B). Alizarin-Red stained individuals were placed on their right side, and the operculum was opened and held with pins to expose the branchial arches. Branchial traits were measured under a Wild M3C stereomicroscope (Wild Leitz Ltd, Heerbrugg, Switzerland) by a single observer. Gill raker number was defined as the number of major Alizarin Red-positive rakers on the ceratobranchial and epibranchial gill arch bones. Gill raker length was measured on the second major raker on the ceratobranchial, counting from the epibranchial-ceratobranchial joint. Gill arch length was measured from this joint to the anterior end of the ceratobranchial.

### Testing for inversion genotype–trait associations and effect sizes

Only adult individuals (i.e., standard length ≥ 30 mm) were included in the analyses. Trait values were classified as outliers using the boxplot method with the function *is_extreme* (Kassambara, 2026). Briefly, any value below the first quartile (Q1) minus three times the interquartile range or above the third quartile (Q3) plus the interquartile range (IQR = Q3-Q1) were removed from further analyses. Final sample sizes for each trait and dataset are provided in Tables S1 and S2. All analyses and plotting were conducted in R version 4.4.3 (R Core Team, 2025).

If inversions contribute to ecotype divergence, the largest phenotypic differences are expected between individuals homozygous for the marine (MAR) and freshwater (FW) inversion orientations. Thus, traits differing between inversion homozygotes were identified using a mapping approach based on Archambeault et al. (2020). This approach compares two models to determine whether inversion genotype explains trait variation: a “null model” containing only covariates and an “alternative model” containing the same covariates plus genotype at the focal inversion.

To define model covariates and response distributions, models were fitted using the package *glmmTMB* (McGillycuddy et al., 2025). Standard length and sex were tested as covariates for all traits. Staining quality was additionally tested for all lateral line neuromast traits, and lateral plate number was tested for neuromast number in the main trunk posterior line (Mp), as these factors are known to influence neuromast number in this line (Wark et al., 2012).

Additional covariates were tested depending on the dataset. For the lab-raised datasets, the effect of housing-tank was tested on all traits. Cross was not included as an additional covariate because cross and housing tank largely overlapped, with individuals from the same cross housed together in one or two tanks. Thus, accounting for possible variation among housing tanks also accounts for much of the variation among crosses. Photographs of the lab-raised marine individuals were taken by three different observers, and observer was therefore tested as covariate for linear body shape traits. Observer was not tested for external count traits because each housing-tank was only measured by one observer, such that housing tank also captures possible observer effects. All relevant covariates described above were initially included in a model, and their effects were tested using the *drop1* function. A covariate was kept for the final model when its *p*-value was below 0.05.

Model assumptions were assessed by simulating and plotting residuals from the fitted null models using the DHARMa package (Hartig, 2024). Models were fitted using a Gaussian response distribution, except for a few count traits that required a generalized Poisson distribution. Model covariates and response distributions used for trait mapping are described in Table S4.

To test the effect of inversion genotype, null and alternative models were compared using a Chi-square test implemented with the function *anova.* The resulting *p*-values were used to assess the statistical support for an association between inversion genotype and trait variation. To account for multiple testing, *p*-values were adjusted using the Benjamini–Hochberg False Discovery Rate (FDR) procedure (Benjamini & Hochberg, 1995), implemented with the function *p.adjust*. Associations with FDR-adjusted *p*-value < 0.05 were considered statistically strongly supported.

The effect strength of inversion genotype was calculating as the percent phenotypic variation explained (PVE), derived from the log-of-odds (LOD) score of the model comparison. An approximation of the LOD score was calculated to allow the same approach to be applied across different types of models. Specifically, this approach estimates the log-likelihood (logLik) for the null (m0) and alternative (m1) models using the function *logLik*, which are then used to calculate the following LOD approximation:

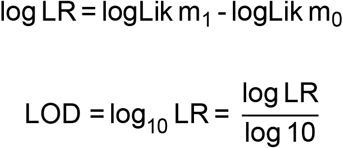

The LOD score was used to calculate the PVE using the following formula (Broman & Sen, 2009):

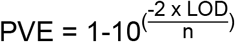

Trait-to-inversion mapping was performed for 33 traits: 17 linear body shape traits (Fig. S1A, Table S3), three branchial traits (Fig. S1B), 12 lateral line neuromast traits (Fig. S1C), and lateral plate number. For the wild-caught dataset, the association between inversion genotype and lateral plate number was tested separately for low-plated fish (≤ 10 lateral plates) and fish with more than 10 lateral plates (so 34 traits). Analyses were conducted independently for the wild-caught stream, lab-raised stream, and lab-raised marine datasets.

### Consistency of inversion–trait associations among datasets

To compare results between the wild-caught stream, lab-raised stream, and lab-raised marine datasets, a list of inversion–trait associations with strong or marginal statistical support was compiled for each dataset. This included seven associations with strong statistical support (FDR-adjusted *p*-value < 0.05) for each of the wild-caught stream and lab-raised stream datasets, and five associations for the lab-raised marine dataset. Four associations from the wild-caught stream dataset and one from the lab-raised stream dataset that showed marginal statistical support (FDR-adjusted *p*-values between 0.050 and 0.055) were also included. Across these associations, 15 unique traits were compared among datasets using Cohen’s *d* effect sizes (Cohen, 1988). Lateral plate number was compared only between the wild-caught stream and lab-raised stream datasets because the chrXXI inversion effect was specific to fish with few lateral plates, which were not found in the lab-raised marine individuals. Effect sizes and their 95% confidence interval were approximated using the function *z_to_d* (Ben-Shachar et al., 2020), which converts *z* statistics from any type of glmmTMB model to an approximate Cohen’s *d* value. It was used because standard Cohen’s *d* calculations are not possible for generalized Poisson models.

The direction of effect for each trait was compared among datasets and qualitatively classified based on Cohen’s *d* estimates and their 95% confidence intervals. Inversion–trait associations were classified as “consistent” when the estimates and confidence in at least two datasets were in the same direction and did not overlap with zero. Cases with an opposite Cohen’s *d* estimate sign and confidence intervals were classified as “opposite”.

Estimates with confidence intervals spanning zero were considered to provide insufficient evidence for a directional effect of inversion genotype. Thus, cases when the Cohen’s *d* confidence interval of only one dataset differed from zero were classified as “single dataset”.

### Dominance coefficients and phenotypic effects in heterozygotes

Because clear inversion–trait associations were found only for the chrXXI inversion (see Results), phenotypic effects in heterozygotes were examined only for this inversion. These analyses were conducted in the two Atlantic-derived stream population datasets (i.e., wild-caught stream and lab-raised stream), allowing direct comparison of heterozygote effects expressed in the wild and under standardized laboratory conditions. These analyses included traits for which the Cohen’s *d* estimate and confidence interval did not overlap with zero because it indicates a directional effect of inversion genotype: 12 traits for the wild-caught stream dataset and 11 traits for the lab-raised stream dataset. For these traits the dominance coefficient, *h*, was calculated using the formula *h* = d/a, based on:

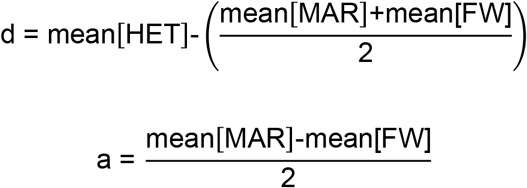

Dominance effects of the marine-orientation, assumed to be the inverted orientation, were classified per trait and dataset following Miller et al. (2014): < -1.25 = underdominant; -1.25 to -0.75 = recessive; -0.75 to -0.25 = partially recessive; -0.25 to 0.25 = additive; 0.25 to 0.75 = partially dominant; 0.75 to 1.25 = dominant; and > 1.25 = overdominant. Mean values of the homozygotes and heterozygotes were estimated from the model including covariates and inversion genotype using the function *emmeans* (Lenth, 2025). When the dominance coefficient indicated overdominance, pairwise comparisons among genotypes were performed using the function *pairs.emmGrid* (Lenth, 2025).

## Results

### The chrXXI, but not the chrI and chrXI, inversion affects multiple traits

In the naturally polymorphic wild-caught stream dataset, there is little statistical support for associations between chrI or chrXI inversion genotype and any of the measured traits (Table S4). In contrast, chrXXI inversion genotype is associated with variation in multiple traits: seven of the 33 measured traits show a statistically well-supported association with inversion genotype (FDR adjusted *p*-value < 0.05) and four additional traits show marginal statistical support (FDR adjusted *p*-value 0.05–0.055; Table 1, Fig. S2). These eleven traits include lateral plate number, five linear body shape traits, and five lateral line neuromast traits.

**Table 1.** The chrXXI inversion has small-to-moderate effects across multiple traits. Traits showing strong or marginal statistical support for an association with the chrXXI inversion genotype after FDR adjustment of *p*-values. Results are from comparison of a “null model” including only the covariates with an “alternative model” including covariates as well as the chrXXI inversion genotype. For each trait, the χ² statistic and corresponding *p*-value from the model comparison, the FDR-adjusted *p*-value, the logarithm of the odds (LOD) score, the percent variance explained (PVE) by inversion genotype, and the number of individuals used in the model (n) are reported.

| Dataset | Trait | $\chi^2$ | $p$-value | FDR adjusted<br>$p$-value | LOD | PVE | n |
| --- | --- | --- | --- | --- | --- | --- | --- |
| Wild-caught stream | Anal fin length | 7.798 | 0.005 | 0.026 | 1.69 | 3.15 | 244 |
|  | Dorsal fin length | 5.617 | 0.018 | 0.055 | 1.22 | 2.25 | 247 |
|  | Pelvic spine length | 5.757 | 0.016 | 0.055 | 1.25 | 2.35 | 242 |
|  | Second dorsal spine length | 5.713 | 0.017 | 0.055 | 1.24 | 2.30 | 245 |
|  | Snout length | 7.735 | 0.005 | 0.026 | 1.68 | 3.12 | 244 |
|  | Ethmoid (ET) neuromast number | 12.546 | 0.000 | 0.004 | 2.72 | 6.11 | 199 |
|  | Infraorbital (IO) neuromast number | 16.894 | 0.000 | 0.001 | 3.67 | 8.22 | 197 |
|  | Main trunk posterior (Mp) neuromast number | 19.194 | 0.000 | 0.000 | 4.17 | 9.42 | 194 |
|  | Mandibular (MD) neuromast number | 11.301 | 0.001 | 0.007 | 2.45 | 5.49 | 200 |
|  | Otic (OT) neuromast number | 6.127 | 0.013 | 0.055 | 1.33 | 3.08 | 196 |
|  | Lateral plate number in low-plated fish | 10.404 | 0.001 | 0.009 | 2.26 | 8.95 | 111 |
| Lab-raised stream | Jaw length | 9.928 | 0.002 | 0.008 | 2.16 | 8.94 | 106 |
|  | Standard length | 12.175 | < 0.001 | 0.004 | 2.64 | 10.76 | 107 |
|  | Ethmoid (ET) neuromast number | 11.710 | 0.001 | 0.004 | 2.54 | 10.46 | 106 |
|  | Infraorbital (IO) neuromast number | 15.898 | < 0.001 | 0.001 | 3.45 | 13.93 | 106 |
|  | Main trunk posterior (Mp) neuromast number | 43.687 | < 0.001 | < 0.001 | 9.49 | 34.04 | 105 |
|  | Mandibular (MD) neuromast number | 15.844 | < 0.001 | 0.001 | 3.44 | 13.88 | 106 |
|  | Oral (OR) neuromast number | 6.073 | 0.014 | 0.051 | 1.32 | 5.57 | 106 |
|  | Otic (OT) neuromast number | 9.092 | 0.003 | 0.011 | 1.97 | 8.29 | 105 |
| Lab-raised marine | Lower peduncle length | 10.285 | 0.001 | 0.015 | 2.23 | 5.01 | 200 |
|  | Infraorbital (IO) neuromast number | 9.272 | 0.002 | 0.019 | 2.01 | 6.64 | 135 |
|  | Main trunk posterior (Mp) neuromast number | 35.901 | < 0.001 | 0.000 | 7.80 | 23.35 | 135 |
|  | Mandibular (MD) neuromast number | 7.791 | 0.005 | 0.035 | 1.69 | 5.65 | 134 |
|  | Oral (OR) neuromast number | 17.368 | < 0.001 | 0.001 | 3.77 | 12.24 | 133 |

Ten of these eleven traits have larger values in homozygotes for the marine orientation (Fig. S2): snout length, dorsal fin length, anal fin length, pelvic spine length, second dorsal spine length, neuromast number in the mandibular (MD), infraorbital (IO), ethmoid (ET), and otic (OT) lateral lines, and lateral plate number. The association between the chrXXI inversion and lateral plate number is only observed in fish with fewer than 10 lateral plates. In contrast to the other lateral line traits, freshwater-orientation homozygotes have more neuromasts in the main trunk posterior (Mp) lateral line than marine-orientation homozygotes.

In the lab-raised stream dataset, associations between chrI and chrXI genotype were not tested, but multiple traits are again associated with chrXXI inversion genotype. Seven inversion–trait associations show strong statistical support (FDR-adjusted *p*-value < 0.05), including two linear body shape traits and five lateral line neuromast traits (Table 1, Fig. S3). Marine-orientation homozygotes have higher values for jaw and standard length, as well as more neuromasts in the MD, IO, ET, and OT lateral lines. In contrast, freshwater-orientation homozygotes have more neuromasts in the Mp lateral line. One additional trait, the OR lateral line, shows marginal statistical support (FDR-adjusted *p*-value = 0.051) with a trend for marine-orientation homozygotes to have larger trait values.

In the lab-raised marine dataset, there is again little statistical support for associations of the chrI and chrXI inversions with the measured traits (Table S4), whereas chrXXI inversion genotype is associated with variation in five traits (Table 1; Fig. S4). Marine-orientation homozygotes have more neuromasts in the MD and IO lateral lines. In contrast, freshwater-orientation homozygotes have more neuromasts in the OR and Mp lateral lines, as well as longer lower peduncles than marine-orientation homozygotes.

### Effects of the chrXXI inversion on individual traits are mostly small

Across datasets, chrXXI inversion genotype generally explains a relatively small percent of variation in individual traits (Table 1). In the wild-caught dataset, PVE is below 10% for all eleven traits associated with chrXXI inversion genotype (PVE = 2.25–9.42). Effects are somewhat larger in the lab-raised datasets but remain below 20% for most traits. In the lab-raised stream dataset, PVE ranges from 5.57 to 13.88% for all traits except Mp neuromast number, for which chrXXI inversion genotype explains 34.04% of the variation. Similarly, in the lab-raised marine dataset, PVE ranges from 5.01 to 12.24% for traits other than Mp neuromast number, for which PVE is 23.35%. Thus, the chrXXI inversion is associated with relatively small amounts of variation in most individual traits, with its strongest effect consistently observed for Mp neuromast number.

### ChrXXI inversion–trait associations are largely consistent across datasets

A total of 15 traits received strong statistical support (FDR *p*-value < 0.05) for an effect of the chrXXI inversion in at least one of the three datasets (Table 1). For nine of these traits, effects are consistent across at least two datasets, with Cohen’s *d* estimates in the same direction and confidence intervals not overlapping zero (Fig. 1, Table 2). Six of these nine consistent associations also had effect estimates in the same direction in all datasets, including the length of the lower peduncle, as well as the neuromast number in four lateral lines on the head (ET, MD, IO, OT) and in the Mp lateral line. The remaining three traits show consistent effect between two datasets: for the second dorsal and pelvic spines the effect estimates of the wild-caught stream and lab-raised marine were in the same direction, while for snout length the effect estimates were in the same direction in the wild-caught and lab-raised stream individuals.

**Figure 1.**
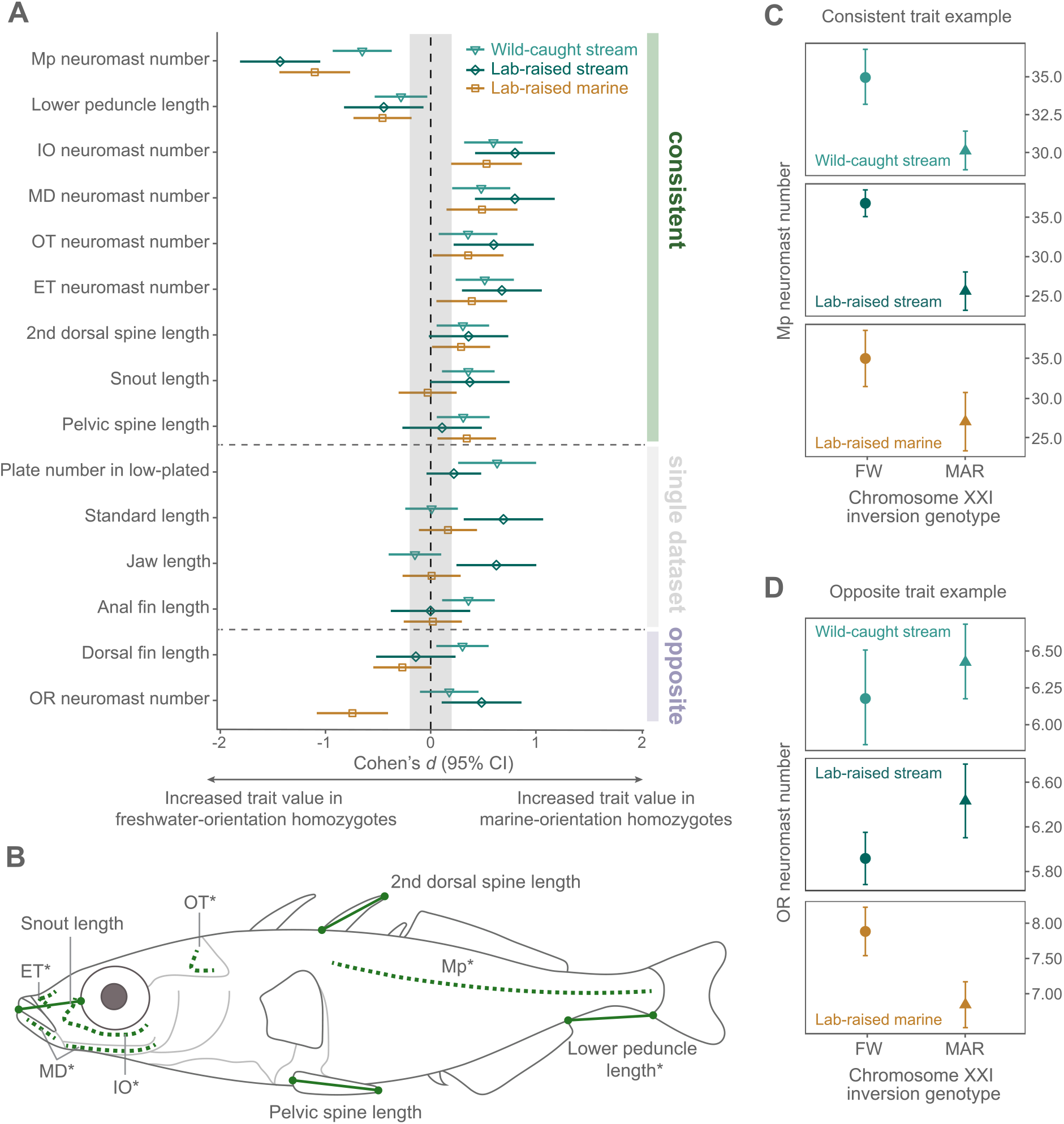
Phenotypic effects of the chromosome XXI inversion. Traits mapped to the chromosome XXI inversion in the wild-caught and lab-raised datasets. (A) Effect of alternative homozygous inversion genotypes on the mapped traits across datasets. Effect sizes are shown as Cohen’s *d* with associated 95% confidence intervals (CI). Positive Cohen’s *d* values indicate increased trait values in homozygotes for the marine orientation, whereas negative values indicate increased trait values in homozygotes for the freshwater orientation. Traits were categorized as either “consistent” (supported effect in the same direction in at least two datasets), “single dataset” (supported effect in only one dataset), or “opposite” (supported effect in opposite direction across datasets) based on the Cohen’s *d* estimates and CI (see Methods and Table 2). Lateral plate number is shown only for the wild-caught and lab-raised stream datasets because the effect is specific to low-plated fish (< 10 lateral plates), which are not found in the lab-raised marine fish. (B) Schematic representation of traits showing consistent effects in at least two datasets. Asterisks indicate traits showing consistent effects across all three datasets. (C) Example of a “consistent” trait: main trunk posterior (Mp) neuromast number. (D) Example of an “opposite” trait: Oral (OR) neuromast number. Panels (C) and (D) show estimated marginal means and 95% CI from the models used to test inversion–phenotype associations on the lab-raised and wild-caught datasets. FW denotes individuals homozygous for the freshwater orientation (filled circles) and MAR denotes individuals homozygous for the marine orientation (filled triangles).

**Table 2.**
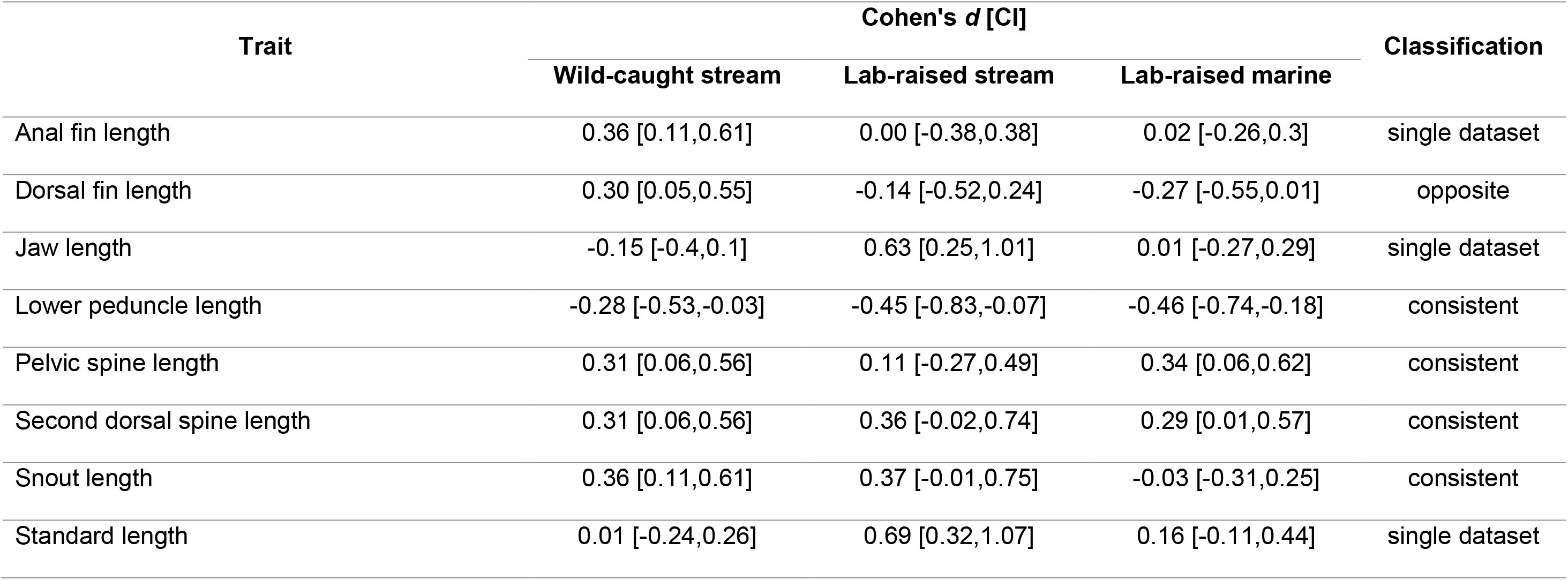
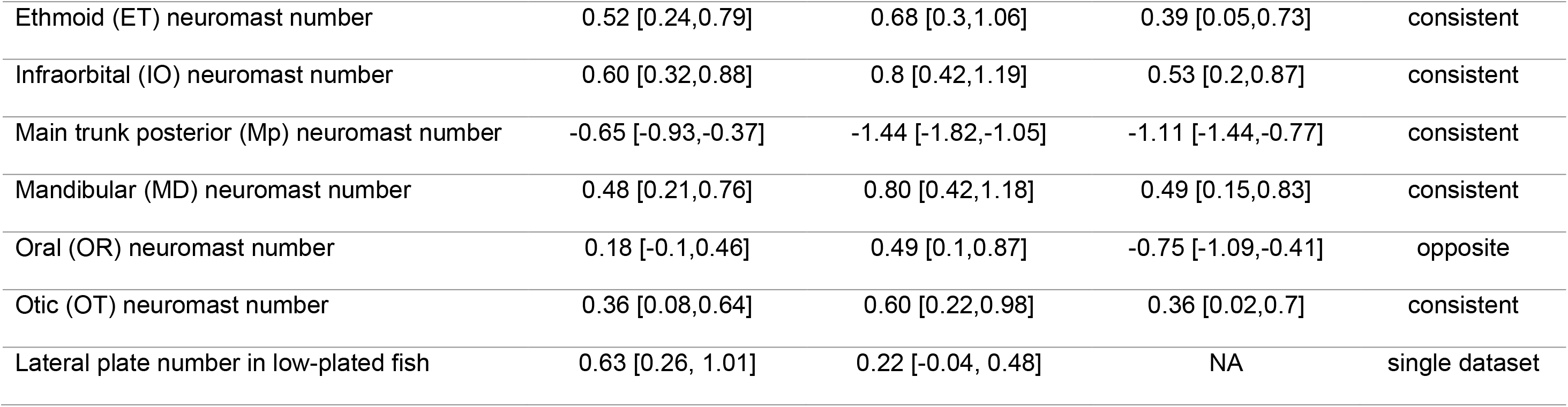
The chrXXI inversion has both consistent and dataset-specific effects on traits. Comparison of Cohen’s *d* estimates for 15 inversion–phenotype associations that received strong statistical support in at least one of the three datasets (see Table 1). For each trait, the Cohen’s *d* estimate and its 95% confidence interval (CI) are reported, together with the classification based on those values (Fig. 1). Positive Cohen’s *d* values indicate that marine-orientation homozygotes have a larger trait value, while negative values indicate that freshwater-orientation homozygotes have a larger trait value. Lateral plate number was not analyzed (NA) in the lab-raised marine dataset because the chrXXI inversion association was specific to low-plated fish (< 10 lateral plates), which were not observed in this dataset. Confidence intervals spanning zero were interpreted as a lack of evidence for an effect of inversion genotype.

Four associations are specific to a single dataset. These include jaw and standard length in the lab-raised stream dataset, and anal fin length and lateral plate number in fish with fewer than 10 plates in the wild-caught dataset. For lateral plate number, the effect estimate in the lab-raised stream dataset is in the same direction, although statistical support for an association is weak (uncorrected *p*-value = 0.46). Finally, two traits show effects in opposite directions between datasets: dorsal fin length and neuromast number in the oral (OR) line (Fig. 1D). In both cases, the opposite effects occur between datasets derived from different geographic origins (eastern Pacific marine versus Atlantic-derived stream populations).

### ChrXXI inversion heterozygotes generally show intermediate phenotypes

In the wild-caught and lab-raised stream datasets, chrXXI heterozygotes are intermediate in almost all traits (Fig. 2). Two traits deviate from this pattern: wild-caught heterozygotes have fewer Mp neuromasts than either homozygote, whereas lab-raised stream heterozygotes have longer second dorsal spines than either homozygote. However, in both cases the estimated mean for heterozygotes is statistically similar to that of marine-orientation homozygotes (Tukey post hoc comparisons: Mp neuromast number, *p*-value = 0.306; second dorsal spine length, *p*-value = 0.862).

**Figure 2.**
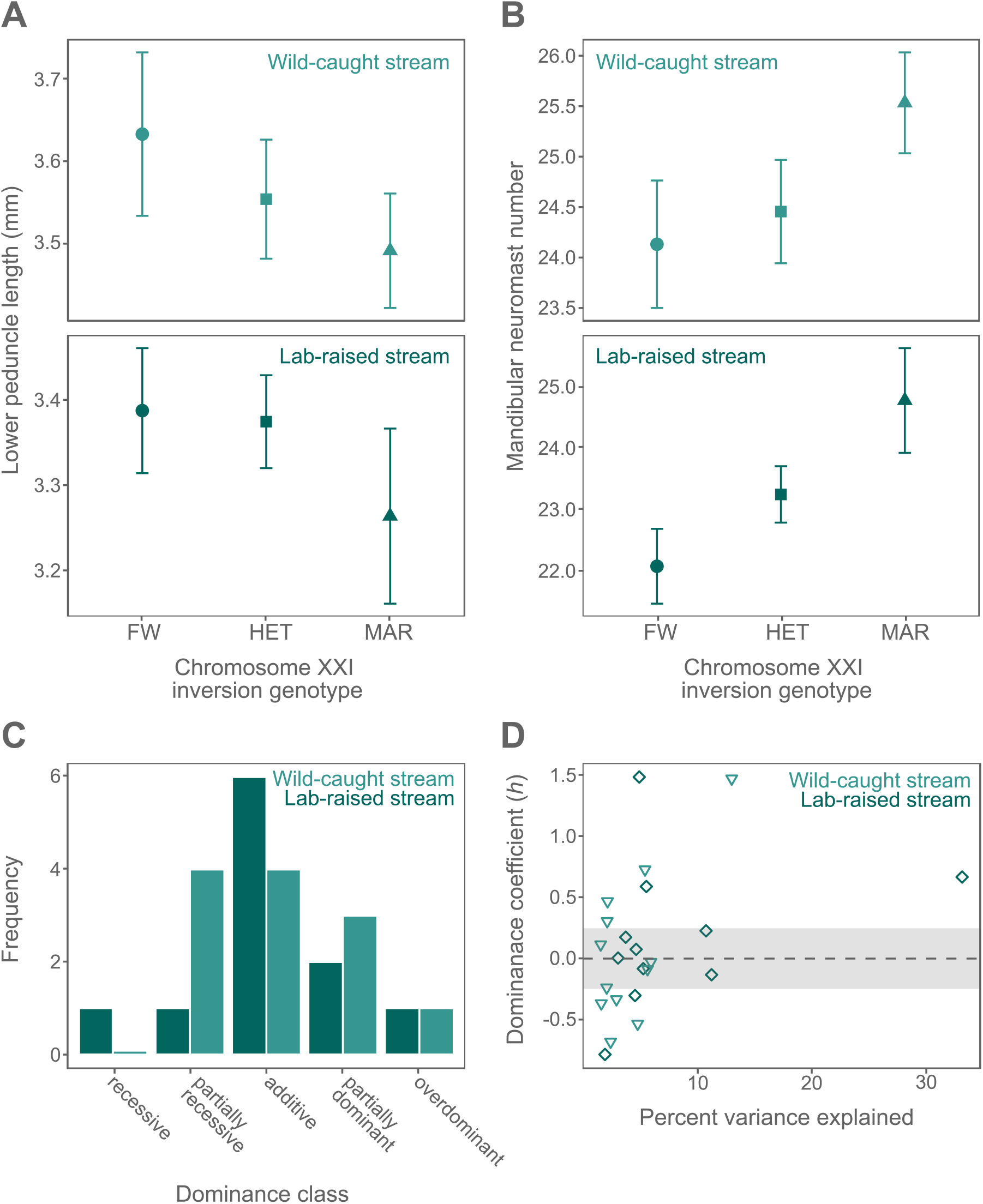
Heterozygotes are phenotypically intermediate across the mapped traits. (A, B) Examples of traits with mostly intermediate trait values in heterozygotes: (A) lower peduncle length and (B) mandibular neuromast number in the wild-caught stream and lab-raised stream datasets. (C) Distribution of dominance classes of the marine-orientation for the lab-raised and wild-caught stream datasets for the traits influenced by the chrXXI inversion. Dominance coefficients (*h*) were calculated as *h* = d/a and categorized into dominance classes as described in Miller et al. (2014). Briefly, values between -0.25 and 0.25 were classified as additive, with more positive values indicating increasing dominance and increasingly negative values indicating increasing recessivity. (D) Dominance coefficients tend towards additive and dominant classes with increasing effect size (percent variance explained). Wild-caught stream data are indicated in light green inverted triangles, and lab-raised stream data are indicated in darker green diamonds.

On average, dominance coefficients of the marine orientation are largely additive in both the wild-caught (*h* = 0.069 ± 0.61) and lab-raised stream datasets (*h* = 0.176 ± 0.59), although individual estimates range from recessive to overdominant (Fig. 2C, Table 3). Traits with higher PVE end to have additive or partially dominant effects (Fig. 2D). Of the eight traits that have an association with the chrXXI inversion in both datasets, only one had the same dominance class in both wild-caught and lab-raised stream individuals. In most cases, however, classifications differed by only one dominance class (i.e., additive versus partial dominance). The exception is the length of the lower peduncle where the marine-orientation is additive in the wild-caught stream dataset and recessive in the lab-raised stream dataset

**Table 3.** Dominance coefficients of the chrXXI inversion indicate largely additive effects. Twelve traits from the wild-caught stream and eleven traits from the lab-raised stream dataset identified in the comparison of homozygote genotypes were used to evaluate the phenotype of heterozygotes. For each trait, the dominance coefficient (*h*) describes the phenotype of heterozygotes relative to the two homozygous genotypes. A positive *h*-value indicates heterozygotes are phenotypically closer to marine-orientation homozygotes, while a negative *h*-value indicates heterozygotes are phenotypically closer to freshwater-orientation homozygotes. Dominance coefficients were calculated and categorized following Miller et al. (2014) and reported with the number of individuals (n) used to calculate *h*. NA indicates traits for which no effect of the chrXXI inversion was identified in the comparison of homozygote genotypes

| Trait | Wild-caught stream |  |  | Lab-raised stream |  |  |
| --- | --- | --- | --- | --- | --- | --- |
| | $h$ | $n$ | Dominance class | $h$ | $n$ | Dominance class |
| Anal fin length | -0.336 | 389 | partially recessive | NA | NA | NA |
| Dorsal fin length | -0.370 | 393 | partially recessive | NA | NA | NA |
| Jaw length | NA | NA | NA | -0.305 | 232 | partially recessive |
| Lower peduncle length | 0.116 | 389 | additive | -0.791 | 232 | recessive |
| Pelvic spine length | 0.308 | 383 | partially dominant | NA | NA | NA |
| Second dorsal spine length | -0.239 | 391 | additive | 1.497 | 233 | overdominant |
| Snout length | 0.472 | 389 | partially dominant | 0.176 | 232 | additive |
| Standard length | NA | NA | NA | -0.083 | 233 | additive |
| Ethmoid (ET) neuromast number | 0.734 | 315 | partially dominant | 0.076 | 226 | additive |
| Infraorbital (IO) neuromast number | -0.087 | 313 | additive | 0.229 | 227 | additive |
| Main trunk posterior (Mp) neuromast number | 1.484 | 307 | overdominant | 0.673 | 227 | partially dominant |
| Mandibular (MD) neuromast number | -0.538 | 317 | partially recessive | -0.133 | 225 | additive |
| Oral (OR) neuromast number | NA | NA | NA | 0.594 | 227 | partially dominant |
| Otic (OT) neuromast number | -0.686 | 312 | partially recessive | 0.004 | 226 | additive |
| Lateral plate number in low-plated fish | -0.029 | 180 | additive | NA | NA | NA |

## Discussion

Chromosomal inversions are thought to play an important role in local adaptation with gene flow, yet key predictions about their phenotypic effects remain largely untested. Here, we investigated inversion–phenotype associations for three divergent chromosomal inversions in threespine stickleback fish, focusing on the number, magnitude, consistency, and dominance of their phenotypic effects. Combining association mapping in a wild polymorphic population with standardized laboratory-raised crosses representing different genomic backgrounds, we found that a 1.7 Mb inversion on chrXXI concentrates variation in multiple traits related to mechanosensation, defense, and swimming. In contrast, we found no evidence for phenotypic associations with the chrI and chrXI inversions. Effects of the chrXXI inversion on individual traits were mostly small to moderate, consistent with the expectation that inversions can combine alleles with relatively small effects on individual traits, thereby contributing to polygenic adaptation through their effect on suppressing recombination.

Heterozygotes generally showed intermediate phenotypes, indicating largely additive effects of the alternative inversion orientation that may allow their frequencies to respond rapidly to divergent selection across habitats. We also found several inversion–trait associations to be shared across different datasets and thus divergent genomic backgrounds, suggesting that consistent genetic differences between the marine and freshwater inversion orientations contribute to parallel phenotypic adaptation across the stickleback distribution. Interestingly, however, some trait associations differed among datasets, suggesting that genomic background and environmental or experimental context can modify inversion-associated phenotypic effects. Overall, our results support the idea that inversions can promote local adaptation by linking co-selected genetic variation underlying multiple ecologically divergent traits.

### Ecological trait divergence is concentrated in the chrXXI inversion

We found that the chrXXI inversion affects multiple and diverse traits, with inversion genotype–phenotype associations involving 15 traits across the three datasets. These traits have been proposed to mediate predator defense, swimming and feeding performance, and detection of mechanical stimuli. For example, variation in caudal peduncle length, which is the length of the narrow posterior region connecting the body to the tail fin (Fig. 1), is associated with different swimming modes, where shorter caudal peduncles increase fast-start swimming performance (Blake, 1983). Fast-start swimming is key for escaping predators, while two other traits found to be associated with the chrXXI inversion (i.e. longer spines and more lateral bony plates) can increase fish survival during handling attempts from vertebrate predators (Lescak & von Hippel, 2011; Reimchen, 1983, 2000). Thus, sticklebacks carrying the marine orientation of the chrXXI inversion may perform better in environments with increased vertebrate predation through the combined defensive function of shorter caudal peduncles, longer spines, and more lateral plates. Beyond predator defense, the inversion-associated changes in snout length and in the number of sensory neuromasts of the lateral line could influence feeding. The longer snouts of marine-orientation homozygotes parallel the longer snouts found in zooplankton feeders (Bentzen & McPhail, 1984; Liem, 1993; McPhail, 1984), while the increased number of head neuromasts may enhance the detection of small planktonic prey, as observed in gobies (Nickles et al., 2020). On the other hand, an increase in neuromasts along the main trunk can facilitate spatial navigation (Jiang et al., 2017; Montgomery et al., 2014). In line with this prediction, sticklebacks living in complex habitats, such as stream and benthic fish, have more neuromasts along the main trunk Mp line compared to marine and limnetic fish (Wark & Peichel, 2010). Consistent with this pattern, the chrXXI freshwater homozygotes have more neuromasts in the Mp line. Together, these patterns provide evidence that the chrXXI inversion influences multiple traits with diverse ecological functions, often in the direction that is consistent with phenotypic divergence among natural stickleback ecotypes.

The concentration of effects on diverse traits within the chrXXI inversion is particularly notable given its relatively small size: the inversion only contains about 85 genes and spans 1.7 Mb of the approximate 450 Mb stickleback genome. Thus, the number of different traits associated with the chrXXI inversion exceeds what would be expected based on its genomic size alone but instead suggests an unusually dense concentration of phenotypic divergence within that genome region. Indeed, many inversions associated with multiple traits in other systems are substantially larger and, in some cases, span almost entire chromosomes (Hager et al., 2022; Kapun et al., 2016; Mérot et al., 2020; Sanchez-Donoso et al., 2022; Wellenreuther et al., 2025). Overall, the notable concentration of ecologically relevant traits within the chrXXI inversion is consistent with the local adaptation mechanism proposed by Kirkpatrick and Barton (2006), in which inversions are adaptive because they capture multiple adaptive alleles that are favored through selection on their individual fitness effects. Although we cannot rule out that some of the observed effects stem from the same pleiotropic locus, the diversity of the affected traits is consistent with contributions from multiple alleles. Thus, we propose that the association of multiple traits at least partly reflects linkage mediated by the chrXXI inversion, with the alternative orientations favored by combined selection on multiple alleles affecting ecologically relevant traits.

### The chrXXI inversion concentrates small effects as predicted to prevent swamping under gene flow

Our results also show that the chrXXI inversion has small-to-moderate effects on individual traits, which are likely to arise from multiple small-effect alleles. Such alleles are prone to being “swamped” or lost under high gene flow and are thus individually unlikely to resist homogenizing gene flow and contribute to local adaptation. However, theoretical work shows that local adaptation can evolve with swamping-prone alleles when they are concentrated within a non-recombining genomic region (Flaxman et al., 2014). An inversion can concentrate such alleles and combine their small effects so that the inversion behaves like a large-effect “supergene”. Thus, our findings support the theoretical idea that chromosomal inversions can facilitate local adaptation with gene flow by concentrating multiple small-to-moderate alleles into a single linked genomic region (Roesti, 2018; Schaal et al., 2022). This scenario may be of particular importance for polygenic adaptation, as is the case in sticklebacks (Arnegard et al., 2014; Jones et al., 2012; Laurentino et al., 2020; Reid et al., 2021), for which genetic variation would otherwise be spread across the genome and easily homogenized by gene flow.

### Intermediate phenotypes of heterozygotes may facilitate ecological divergence and persistence in the ancestral gene pool

Heterozygotes at an inversion should be disfavored when alternative inversion orientations are favored in different habitats because they will have lower fitness compared to the locally adapted homozygote. In this scenario, alternative orientations are associated with different phenotypic optima and heterozygotes deviate from it. Here, we show that heterozygotes at the chrXXI inversion have mostly intermediate phenotypes, which could translate into reduced fitness relative to the locally adaptive homozygote in each habitat. Previous studies have found similar patterns for inversions involved in ecological divergence (Akopyan et al., 2026; Hager et al., 2022; Koch et al., 2021; Lowry & Willis, 2010; Sanchez-Donoso et al., 2022). Though phenotypic intermediacy of heterozygotes does not necessarily mean intermediate fitness, intermediate phenotypes are known to be selected against in the benthic–limnetic stickleback ecotypes (Arnegard et al., 2014; Schluter, 1995). Therefore, if alternative inversion orientations are favored due to their divergent phenotypic optima, intermediate heterozygote phenotypes will have reduced fitness in either habitat and thereby favor the spread of the locally adaptive orientation. Nonetheless, testing this prediction will require validating fitness consequences of the phenotypes identified here in both the local and foreign habitats to determine how selection acts on the chrXXI inversion.

Across the identified traits influenced by the chrXXI inversion, its phenotypic effects are largely additive. This observation aligns with a previous QTL mapping study which suggested that a tendency towards additivity could be a general feature in stickleback phenotypic evolution (Miller et al., 2014). This trend was proposed to result from a bias in the mutations favored by selection, given the extensive use of standing genetic variation that has been sorted by selection in both marine and freshwater populations (Barrett & Schluter, 2008; Colosimo et al., 2004; Haenel et al., 2019; Jones et al., 2012; Roesti et al., 2014). However, as dominance effects can be context-dependent, it would be valuable to explore whether the tendency towards additivity is also found in individuals raised in marine habitats. This could be particularly important for understanding how the freshwater orientation of the chrXXI inversion persists in marine populations where they are potentially maladaptive. Marine sticklebacks are thought to maintain a small number of freshwater alleles which can mediate rapid adaptation to newly formed freshwater habitats (Schluter & Conte, 2009).

However, regions of low recombination, such as inversions, may carry multiple freshwater-adaptive alleles and could therefore be expected to be efficiently removed by selection in marine populations. How, then, is the chrXXI inversion maintained? One possibility is that dominance of these freshwater alleles may be recessive in the sea resulting in a mostly neutral locus. This would facilitate its persistence at low frequency, while the largely additive effects in the freshwater environment allows the inversion to spread during freshwater colonization. Future studies could test whether the magnitude and dominance of phenotypic inversion effects are environmentally-dependent and how this may determine the ability of inversions to persist in habitats where they are maladaptive and, ultimately, contribute to parallel evolution (Westram et al., 2022).

### Context-dependent inversion effects may shape population-specific adaptation

We found that some inversion–phenotype associations differed across datasets. While we cannot rule out that some of these differences arose from inherent differences among datasets, such as in sample size or other aspects of study design, our results suggest that inversions can have effects that depend on the biological context. Previous studies have similarly reported phenotypic effects of inversions to differ among populations (Arostegui et al., 2019; Nicolas et al., 2026; Weinstein et al., 2019; but see Kapun et al., 2016; Lowry & Willis, 2010). Such differences could arise from epistatic interactions between the inversion and genetic variation elsewhere in the genome.

The genetic context-dependent effect of the chrXXI inversion on lateral plate number appears to provide such an example. We found that the chrXXI inversion affects lateral plate number only in low-plated fish caught from a natural stream. In contrast, there was no phenotypic effect of the inversion on lateral plate number in completely-plated fish in both wild-caught stream and lab-raised marine individuals. This suggests an interaction between the chrXXI inversion and the genetic basis of plate morph. Consistent with this interpretation, Wark et al. (2012) reported a similar interaction between QTL on chromosomes XXI and IV, the latter containing *Eda,* a major gene controlling plate morph in sticklebacks (Colosimo et al., 2004). Thus, epistasis between the chrXXI inversion and *Eda* on chromosome IV could explain the absence of an inversion effect in completely-plated fish.

Another explanation for differences in inversion–phenotype associations is genetic variation within each inversion orientation. The chrXXI inversion is an old variant (Nelson & Cresko, 2018) and thus expected to have accumulated substantial genetic diversity (Faria et al., 2019). Such variation could generate different haplotypes within the same inversion orientation, potentially allowing populations to fine-tune adaptation to similar but not identical habitats. This may be particularly important because suppressed recombination between orientations can constrain adaptation when novel environments favor new combinations of inversion-linked alleles (Roesti et al., 2022). Even if a population lacks an inversion haplotype carrying the locally optimal variants, genetic redundancy could allow variation elsewhere in the genome to produce similar adaptive phenotypes (Láruson et al., 2020), as appears common in sticklebacks (Poore et al., 2023). Genotype-by-environment interactions could further alter the phenotypic effects of an inversion across environments. Taken together, genetic variation within and outside inversions, together with environmental effects, could generate context-dependent inversion effects and facilitate population-specific adaptation.

### The adaptive role of inversions requires direct testing

A central hypothesis for the adaptive value of inversions is that they link multiple locally adaptive alleles by suppressing recombination (Kirkpatrick & Barton, 2006). Our study directly tested key predictions of this hypothesis by examining the number and magnitude of inversion-associated phenotypic effects across ecologically relevant traits. While we found clear support for multiple phenotypic effects of the chrXXI inversion, we found little evidence for such effects of the chrI and chrXI inversions. For chrI, this lack of association was also observed in an independent naturally inversion-polymorphic population (Appendix S1).

Although both of these inversions show repeated divergence across similar ecological contrasts (Roberts Kingman et al., 2021), they are smaller and contain fewer genes than the chrXXI inversion, perhaps making their apparent adaptive value less likely to arise from linking alleles affecting multiple traits. However, we focused on morphological traits and may have missed relevant phenotypes, including behavioral and physiological effects of these inversions.

More generally, repeated differentiation of an inversion between ecotypes does not demonstrate that selection acts on multiple adaptive alleles captured within it. Inversions can be favored through other mechanisms, including functional effects of their breakpoints (Villoutreix et al., 2021). This may apply to the chrXI inversion, whose breakpoint was hypothesized to alter the structure of a gene, thereby producing putatively different isoforms (Jones et al., 2012). The adaptive basis of the chrI inversion remains less clear: although it contains a candidate gene involved in osmoregulation (Shimada et al., 2011), experimental manipulation of salinity did not support salinity as the primary selective agent acting on the chrI inversion (Rodríguez-Fuentes et al., 2026). Understanding why inversions are ecologically selected therefore requires directly connecting inversion genotype to phenotype, function, and fitness rather than inferring adaptive mechanisms from genomic divergence alone.

## Conclusions

Our findings broadly support the view that inversions aid local adaptation by linking multiple, individually modest phenotypic effects that contribute to divergence across ecological environments. This mechanism could be particularly relevant for polygenic adaptation because swamping-prone alleles can be preserved within an inversion by combining their small effects into a large-effect locus. If such combined phenotypic effect matches the local phenotypic optima, an inversion will be selected thanks to the linkage it creates due to recombination suppression. Selection could even transfer and reuse inversions across large geographical distances given that their phenotypic contributions can be stable in divergent genomic backgrounds. However, inversion effects can also be context-dependent, thereby allowing for population-specific adaptation which highlights the idea that inversions are not monoliths and their effects can be shaped by both the genomic and environmental context. However, direct tests on the role of recombination suppression and population-specific adaptation are still needed. For instance, testing the role of recombination suppression can be achieved by “reversing” inversions to restore recombination between divergent inversion haplotypes and comparing the frequency changes between the reversed and wild-type inversion through time. Such an approach, combined with studies testing inversion–phenotype–fitness associations across diverse geographical areas, will be key to disentangle the evolutionary forces acting on inversion evolution.

## Acknowledgements

We are very grateful to all who contributed with photographs for phenotyping: Eugenio Feliciano, Janine Hunziker, Anna Kropf, Lukas Mosimman, Carmen Sopranis, and Ladina Töndury. We thank Matthias Bopp and Steffen Bader for approving and supporting sampling of the Grasbeuren population, Jelena Jäggi for help in sampling the wild-caught stream populations, R. Benjamin Sulser for support with the automated landmarking pipeline, and Gerard Martínez-De León and Stephan Peischl for support with statistical analysis. Finally, we are grateful to the Peichel lab for feedback during the development of this project. We acknowledge the use of Claude (AI) as a support tool for analysis and visualization scripts. All scientific content, analyses, interpretations, and writing of the manuscript were reviewed and approved by the authors. This project was funded by Swiss National Science Foundation grant 310030_204681 to CLP.

## Conflict of interest statement

The authors declare no conflict of interest.

## Data availability statement

All code and raw data are available in Zenodo at 10.5281/zenodo.22896765

## Research ethics statement

All experimental procedures were performed in accordance with the relevant ethical standards and regulations. Animal husbandry and experiments performed on live animals were approved by the Veterinary Service of the Office of Agriculture and Nature of the Canton of Bern (VTHa-Nr. BE4/16, BE103/2021, BE43/2025).

## Author contributions

Juliana Rodríguez-Fuentes: Data Curation – lead, Formal analysis – lead, Investigation – lead, Methodology – equal, Visualization – lead, Writing – Original Draft – equal. Lynn Wedekind: Investigation – supporting, Writing – Review & Editing – supporting. Lea Stadelmann: Investigation – supporting, Writing – Review & Editing – supporting. Nicole Nesvadba: Investigation – supporting, Writing – Review & Editing – supporting. Verena Saladin: Investigation – supporting, Writing – Review & Editing – supporting. Marius Roesti: Conceptualization – equal, Methodology – equal, Project administration – supporting, Resources – equal, Supervision – supporting, Visualization – supporting, Writing – Original Draft – equal. Catherine L. Peichel: Conceptualization – equal, Funding acquisition – lead, Methodology – equal, Project administration – lead, Resources – equal, Supervision – lead, Writing – Original Draft – equal.

## Notes

https://doi.org/10.5281/zenodo.22896765

### Competing Interest Statement

The authors have declared no competing interest.

