## Supplementary figures and appendix S1 for "A chromosomal inversion acts as a hotspot for divergence in multiple ecologically relevant traits"

**Supplementary material**

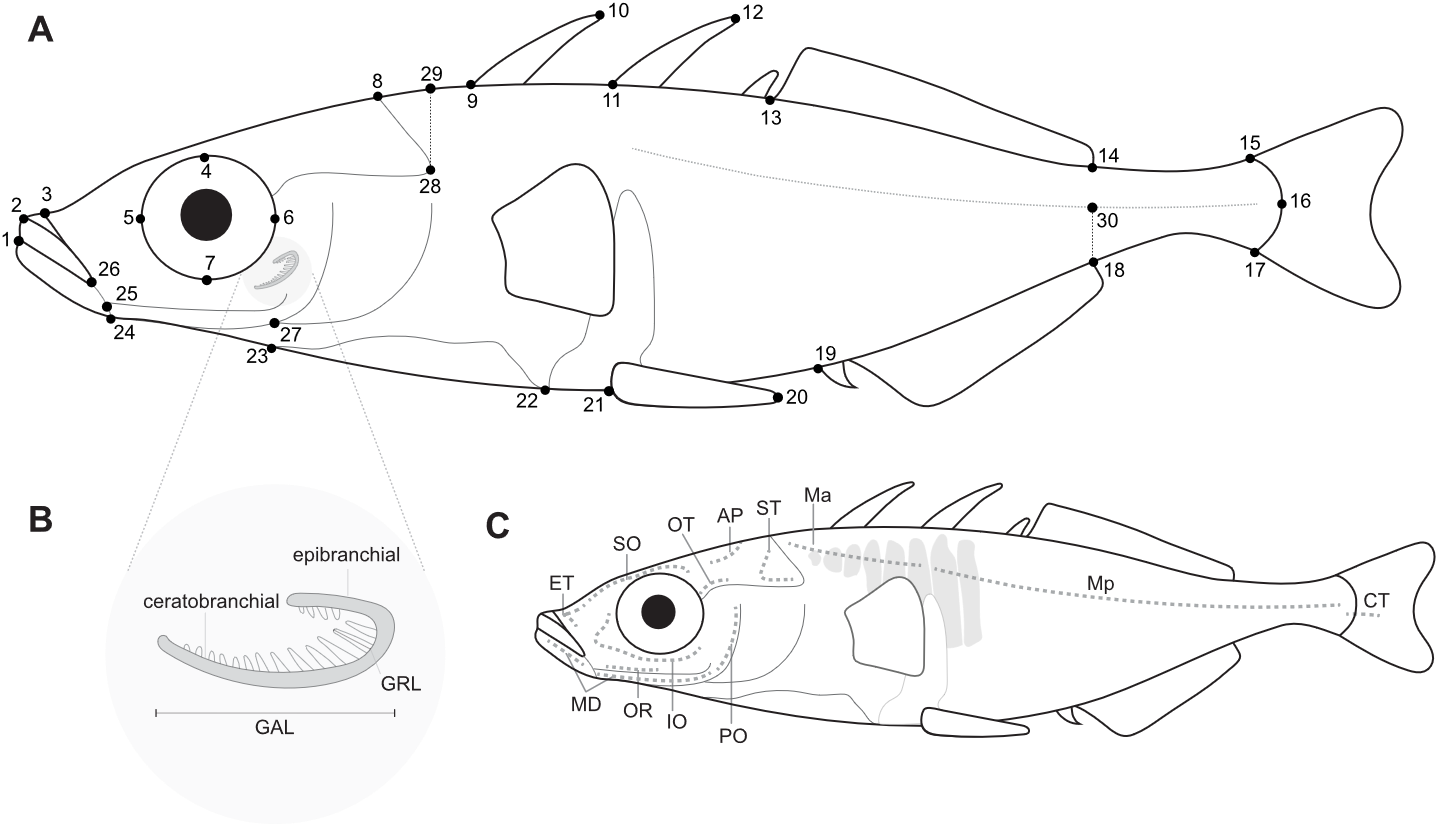

**Figure S1.** **Schematic representation of morphological traits measured.** (A) Landmarks used to calculate the linear body shape traits as described in Table S3. (B) Branchial feeding traits measured on the first branchial arch include: gill arch length (GAL), length of the second gill raker on the ceratobranchial (GRL), and the count of rakers on the ceratobranchial (ventral) and epibranchial (dorsal). (C) External count traits include: neuromast number in the ethmoid (ET), supraorbital (SO), otic (OT), anterior pit (AP), supratemporal (ST), main trunk anterior (Ma), main trunk posterior (Mp), caudal fin (CF), preopercular (PO), infraorbital (IO), oral (OR), and mandibular (MD) lines, as well as lateral plate number.

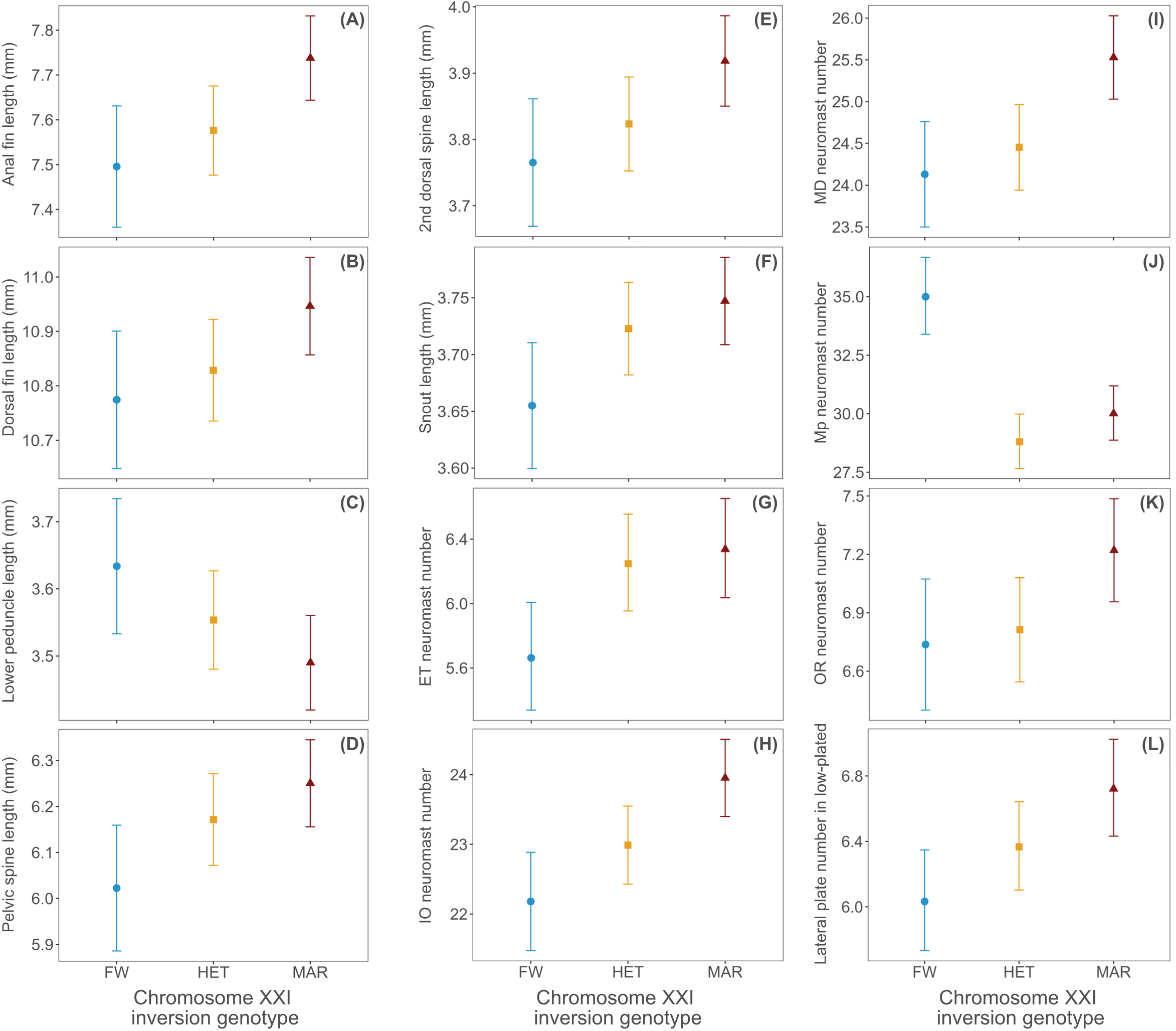

**Figure S2. Significant inversion–phenotype associations in the wild-caught stream GRA population.** Estimated marginal means and 95% confidence intervals for the linear models including inversion genotype and covariates for: the length of the (A) anal fin, (B) dorsal fin, (C) lower peduncle, (D) pelvic spine, (E) second dorsal spine, and (F) snout; neuromast number in the (G) ethmoid (ET), (H) infraorbital (IO), (I) mandibular (MD), (J) main trunk posterior (Mp), and (K) oral (OR) lateral lines; and (L) lateral plate number in low-plated fish. These include 11 traits with strong or marginal statistical support for an effect of inversion genotype (Table 1). One additional trait, lower peduncle length, was included because it has evidence for a directional effect of inversion genotype based on the Cohen’s *d* estimate (Table 2). FW denotes individuals homozygous for the freshwater orientation (blue circles), HET denotes heterozygous individuals (yellow squares), and MAR denotes individuals homozygous for the marine orientation (red triangles).

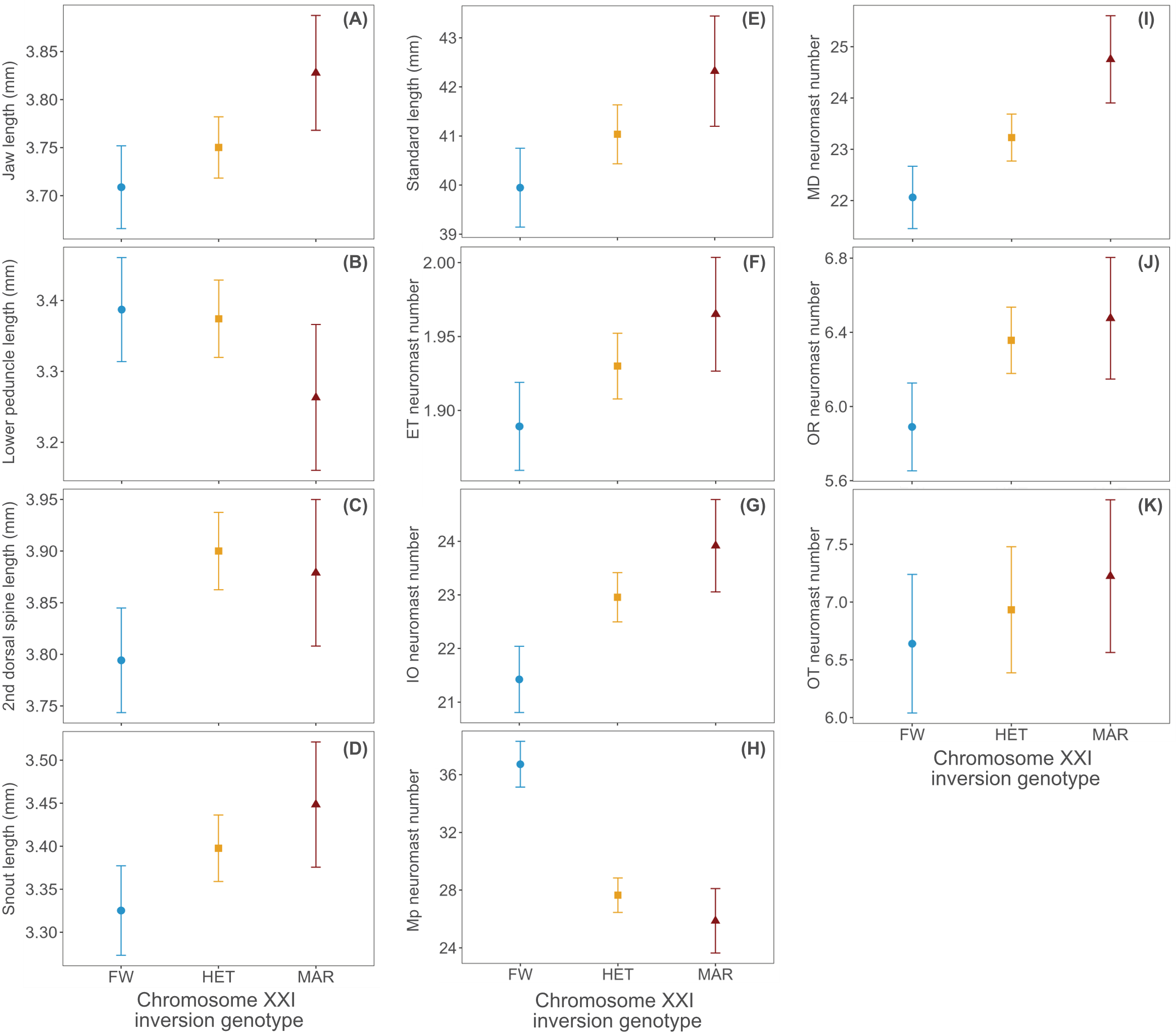

**Figure S3. Significant inversion–phenotype associations in the lab-raised stream dataset.** Estimated marginal means and 95% confidence intervals for the linear models including inversion genotype and covariates for: the length of the (A) jaw, (B) lower peduncle, (C) second dorsal spine, (D) snout, and (E) body (i.e., standard length); neuromast number in the (F) ethmoid (ET), (G) infraorbital (IO), (H) main trunk posterior (Mp), (I) mandibular (MD), (J) oral (OR), and (K) otic (OT) lateral lines. These include eight traits with strong or marginal statistical support of an effect of inversion genotype (Table 1). Three additional traits, lower peduncle, second dorsal spine, and snout length, were included because they have evidence for a directional effect of inversion genotype based on the Cohen’s *d* estimate (Table 2). FW denotes individuals homozygous for the freshwater orientation (blue circles), HET denotes heterozygous individuals (yellow squares), and MAR denotes individuals homozygous for the marine orientation (red triangles).

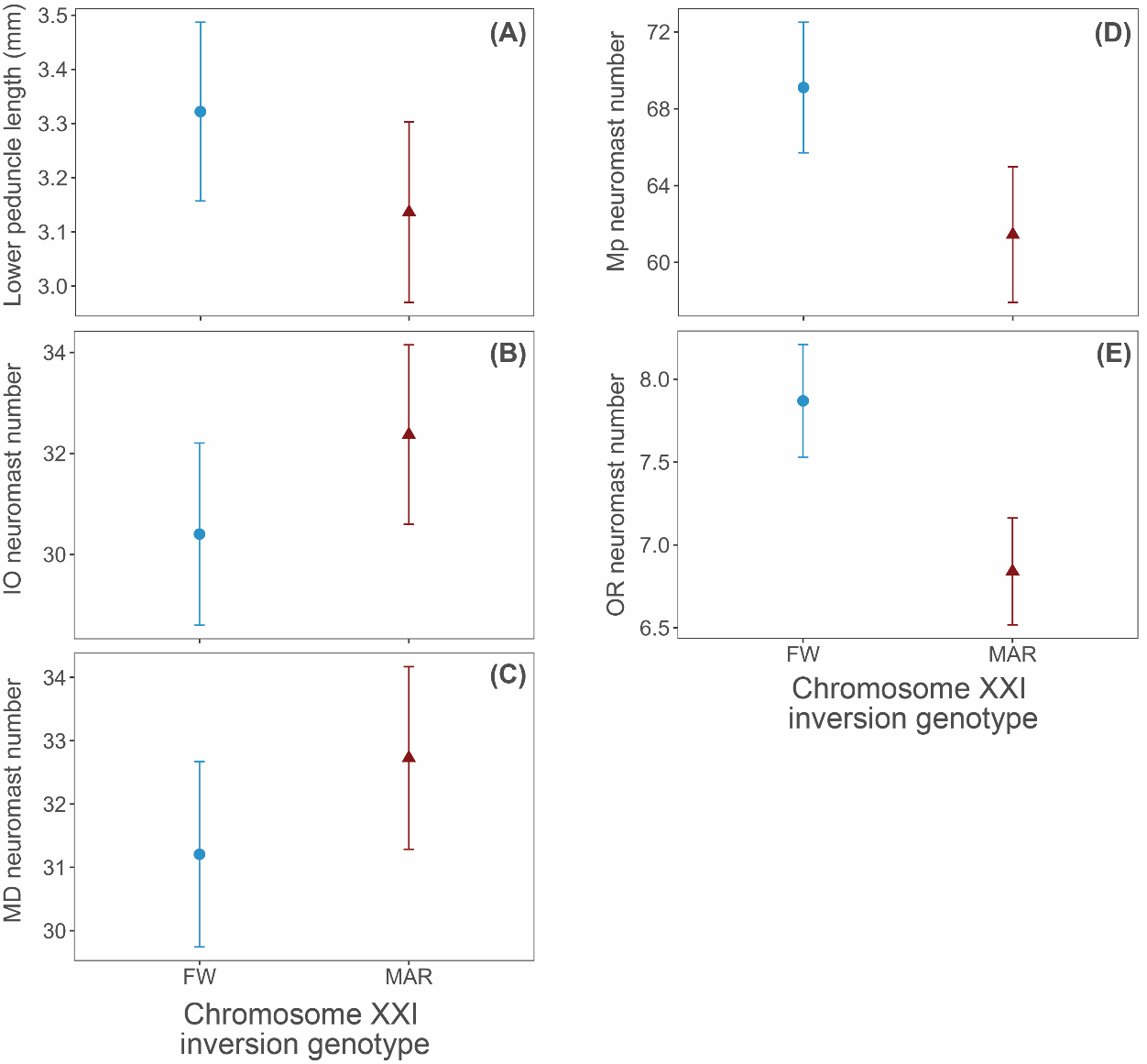

**Figure S4. Significant inversion–phenotype associations in the lab-raised marine dataset.** Estimated marginal means and 95% confidence intervals for the models including chrXXI inversion genotype and covariates for: (A) lower peduncle length; and neuromast number in the (B) infraorbital (IO), (C) mandibular (MD), (D) main trunk posterior (Mp), and (E) oral (OR) lateral lines. These traits have strong or marginal statistical support of an effect of inversion genotype (Table 1). FW denotes individuals homozygous for the freshwater-orientation (blue circles) while MAR denotes individuals homozygous for the marine-orientation (red triangles).

**Appendix S1**

An additional wild population was sampled from the Lake Constance watershed. Wild-caught individuals were collected using minnow traps from the stream population in Salmsacher Aach in Nideraach, Switzerland (NID, 47.54755; 9.20839) during June 2020 (n = 220) and 2021 (n = 14). Sampling was authorized by the Hunting and Fishing Administration of canton Thurgau, under the permits TG-20.04.2020 and TG-15.02.2024. Individuals were processed as described in the Methods. Briefly, a fin clip stored in ethanol was used for DNA extraction as template for PCR to genotype the chrI, XI, and XXI inversions and sex. Bodies were stained with alkaline phosphatase to highlight the neuromasts of the lateral line system. They were additionally stained with Alizarin-Red to highlight bony tissue for linear body shape and branchial feeding traits. All individuals were phenotyped for all traits (Table SI).

This additional stream population (Nideraach, NID) from the same watershed as GRA is only polymorphic at the chrI and chrXXI inversions. The population is fixed for the marine-orientation at the chrXI inversion, and there were no freshwater-orientation homozygotes at the chrXXI inversion. Thus, the phenotypic effects of chromosomal inversions on trait divergence between homozygotes could only be explored for chrI. Association mapping showed that the chrI inversion does not influence any of the measured traits (Table SII).

**Table SI.** **Wild-caught dataset**. Numbers of individuals per sampling season that were phenotyped from the Salmsacher Aach in Nideraach population

| Sampling season | Number of phenotyped individuals | Phenotypes measured |
| --- | --- | --- |
| 2020 | 205 | Branchial feeding, linear body shape, and count traits |
| 2021 | 11 | Branchial feeding, linear body shape, and count traits |

**Table SII. Test of inversion–phenotype associations in the wild-caught NID dataset.** Results from the comparison of a “null-model” including only the covariates versus an “alternative model” including covariates and genotype at the focal inversion for non-significant associations. *P*-values are reported for the model comparison done with ANOVA and its adjusted value after FDR adjustment, as well as the log-of-odds (LOD) score, the percent variance explained (PVE) by the inversion genotype, and the number of individuals used in the model (n). Covariates used in the models to explore the inversion genotype-phenotype associations across the 34 traits measured are listed. Covariate selection depends on the trait category, dataset, and association with standard length. Trait abbreviations follow the names described in Fig. S1. “SL_landmarks” correspond to the standard length of an individual.

| Trait | Covariates | Response distribution | Model comparison *p*-value | X^2^ | FDR adjusted *p*-value | LOD | PVE | n |
| --- | --- | --- | --- | --- | --- | --- | --- | --- |
| Mouth length | SL_landmarks*Sex | Gaussian | 0.889 | 0.020 | 0.950 | 0.004 | 0.018 | 109 |
| Jaw length | SL_landmarks*Sex | Gaussian | 0.853 | 0.034 | 0.950 | 0.007 | 0.032 | 109 |
| Snout length | SL_landmarks*Sex | Gaussian | 0.558 | 0.343 | 0.897 | 0.074 | 0.314 | 109 |
| Epaxial muscle height | SL_landmarks+Sex | Gaussian | 0.415 | 0.665 | 0.897 | 0.144 | 0.608 | 109 |
| Head length upper | SL_landmarks*Sex | Gaussian | 0.373 | 0.794 | 0.897 | 0.172 | 0.726 | 109 |
| Buccal length | SL_landmarks*Sex | Gaussian | 0.479 | 0.501 | 0.897 | 0.109 | 0.458 | 109 |
| Pelvic spine length | SL_landmarks | Gaussian | 0.660 | 0.194 | 0.897 | 0.042 | 0.178 | 109 |
| First dorsal spine length | SL_landmarks | Gaussian | 0.021 | 5.357 | 0.234 | 1.163 | 4.796 | 109 |
| Second dorsal spine length | SL_landmarks | Gaussian | 0.090 | 2.881 | 0.762 | 0.626 | 2.608 | 109 |
| Dorsal fin length | SL_landmarks | Gaussian | 0.531 | 0.392 | 0.897 | 0.085 | 0.359 | 109 |
| Upper peduncle length | SL_landmarks+Sex | Gaussian | 0.809 | 0.059 | 0.950 | 0.013 | 0.054 | 109 |
| Lower peduncle length | SL_landmarks+Sex | Gaussian | 0.922 | 0.010 | 0.950 | 0.002 | 0.009 | 109 |
| Anal fin length | SL_landmarks+Sex | Gaussian | 0.178 | 1.813 | 0.897 | 0.394 | 1.649 | 109 |
| Anterior body depth | SL_landmarks*Sex | Gaussian | 0.255 | 1.298 | 0.897 | 0.282 | 1.183 | 109 |
| Posterior body depth | SL_landmarks+Sex | Gaussian | 0.448 | 0.576 | 0.897 | 0.125 | 0.527 | 109 |
| Eye diameter | SL_landmarks+Sex | Gaussian | 0.504 | 0.447 | 0.897 | 0.097 | 0.409 | 109 |
| Standard length | Sex | Gaussian | 0.992 | 0.000 | 0.992 | 0.000 | 0.000 | 109 |
| Gill raker number (GRN) | SL_landmarks*Sex | Gaussian | 0.008 | 6.997 | 0.203 | 1.519 | 8.276 | 81 |
| Gill raker length (GRL) | SL_landmarks+Sex | Gaussian | 0.659 | 0.195 | 0.897 | 0.042 | 0.244 | 80 |
| Gill arch length (GAL) | SL_landmarks+Sex | Gaussian | 0.610 | 0.260 | 0.897 | 0.056 | 0.329 | 79 |
| Preopercular neuromast number (PO) | Sex | Gaussian | 0.287 | 1.135 | 0.897 | 0.246 | 1.408 | 80 |
| Mandibular neuromast number (MD) | Staining | Gaussian | 0.908 | 0.013 | 0.950 | 0.003 | 0.017 | 80 |
| Oral neuromast number (OR) | Sex | Generalized Poisson | 0.578 | 0.309 | 0.897 | 0.067 | 0.390 | 79 |
| Infraorbital neuromast number (IO) | Sex | Gaussian | 0.572 | 0.319 | 0.897 | 0.069 | 0.408 | 78 |
| Ethmoid neuromast number (ET) | None | Generalized Poisson | 0.742 | 0.109 | 0.934 | 0.024 | 0.147 | 74 |
| Supraorbital neuromast number (SO) | None | Gaussian | 0.893 | 0.018 | 0.950 | 0.004 | 0.025 | 74 |
| Otic neuromast number (OT) | Sex | Generalized Poisson | 0.324 | 0.973 | 0.897 | 0.211 | 1.209 | 80 |
| Anterior pit neuromast number (AP) | None | Generalized Poisson | 0.012 | 6.321 | 0.203 | 1.373 | 8.082 | 75 |
| Supratemporal neuromast number (ST) | None | Generalized Poisson | 0.226 | 1.465 | 0.897 | 0.318 | 1.814 | 80 |
| Main trunk anterior neuromast number (Ma) | SL_landmarks+Sex | Gaussian | 0.536 | 0.383 | 0.897 | 0.083 | 0.484 | 79 |
| Main trunk posterior neuromast number (Mp) | Sex+Plates | Gaussian | 0.626 | 0.238 | 0.897 | 0.052 | 0.305 | 78 |
| Caudal fin neuromast number (CF) | None | Generalized Poisson | 0.729 | 0.120 | 0.934 | 0.026 | 0.154 | 78 |
| Lateral plates in low-plated fish | Sex | Gaussian | 0.229 | 1.446 | 0.897 | 0.314 | 11.351 | 12 |
| Lateral plates in high-plated fish | Sex | Gaussian | 0.221 | 1.499 | 0.897 | 0.325 | 2.245 | 66 |
